# Isolation of muscle stem cells from rat skeletal muscles

**DOI:** 10.1101/690776

**Authors:** Francesca Boscolo Sesillo, Michelle Wong, Amy Cortez, Marianna Alperin

## Abstract

**Background:** Muscle stem cells (MuSCs) are involved in homeostatic maintenance of skeletal muscles and play a central role in muscle regeneration in response to injury. Thus, understanding MuSC autonomous properties is of fundamental importance for studies of muscle degenerative diseases and muscle plasticity. Rat, as an animal model, has been widely used in the skeletal muscle field, however an efficient approach for MuSC isolation through fluorescence-activated cell sorting from rat muscles has never been described. This work aims to develop and validate an effective protocol for MuSC isolation from rat skeletal muscles.

**Methods:** Tibialis anterior, gastrocnemius, diaphragm, and the individual components of the pelvic floor muscle complex (coccygeus, iliocaudalis, and pubocaudalis) were harvested from female rats and digested for isolation of MuSCs. Three protocols, employing different cell surface markers (CD106, CD56, and CD29), were compared for their ability to isolate a pure MuSC population.

**Results:** Cells obtained using the protocol that relies only on VCAM-1 (CD106) as a positive marker showed high expression of Pax7 upon isolation, ability to progress through myogenic lineage while in culture, and complete differentiation in serum deprived conditions. The protocol was further validated in other skeletal muscles proving to be reproducible.

**Conclusions:** CD106 is an efficient marker for reliable isolation of MuSCs from a variety of rat skeletal muscles.

## INTRODUCTION

Muscle stem cells (MuSCs), which reside between the sarcolemma of muscle fibers and the basal lamina, are required for the maintenance of tissue health in adult muscle and proper muscle regeneration in a case of injury (Cheung and Rando, 2013; Keefe et al., 2015; Mauro, 1961). MuSCs exist in a tightly regulated quiescent state (Cheung and Rando, 2013). MuSC activation and proliferation is induced in response to increased mechanical load imposed on a muscle or to muscle injury (Relaix and Zammit, 2012; Tatsumi et al., 2001). Upon activation, MuSCs progress through the myogenic lineage until fusion with damaged myofibers occurs and muscle repair is achieved (Relaix and Zammit, 2012). The activation and differentiation of MuSCs has been extensively studied leading to the identification of sequentially expressed markers, specific to each step of this process. Pax7, a transcription factor exclusively expressed by quiescent and early-activated MuSCs, is required for their functionality both in homeostatic conditions and during regeneration. Indeed, lack of Pax7 expression in MuSCs *in vivo* results in the absence of muscle regeneration following injury (Lepper et al., 2011; Seale et al., 2000; von Maltzahn et al., 2013). Upon activation, expression of MyoD, a transcription factor responsible for cell’s early commitment, promotes MuSC entry into the cell cycle (Cornelison and Wold, 1997). Finally, a downstream effector of MyoD, myogenin, is activated, inducing terminal differentiation of MuSCs, which can now fuse together to form new myofibers or fuse with the existing myofibers.

Studies of autonomous properties of MuSC populations rely mainly on the use of cell sorting. Isolation of MuSCs has been described in mouse, human, pig, and cow (Alexander et al., 2016; Ding et al., 2018; Ding et al., 2017; Liu et al., 2015; Maesner et al., 2016; Uezumi et al., 2016). Published investigations employ a wide array of cell surface proteins as positive markers for MuSC identification and isolation, namely β1-integrin (CD29), CXCR4 (CD184), VCAM-1 (CD106), NCAM (CD56), α-7 integrin, CD34, tetraspanin (CD82), and CD318. Negative selection markers, overall, are conserved among research groups and different mammalian species, and most commonly include CD45 (lymphocytes), CD31 (endothelial cells), CD11b (macrophages), and Sca1 (fibro-adipogenic progenitors). Despite the extensive knowledge of MuSC identification markers, and the broad spectrum of protocols employed for their isolation among multiple species, purification of MuSCs from rat has not been reported to date.

The rat model has been extensively used in skeletal muscle research over the years (Homberg et al., 2017). Rat, compared to other rodent models, better recapitulates human muscle architecture, physiology, and anatomy, making it a better model for the study of skeletal muscles. First, muscle architecture (macroscopic arrangement of muscle fibers), which is fundamental for in vivo muscle function, has been shown to be similar between rats and humans, when compared to other animal models (Lieber and Friden, 2000). Indeed, comparative studies of abdominal muscle architecture revealed a high degree of similarity within the same muscle groups between rat and human. The major architectural parameters, including physiological cross sectional area, operational sarcomere length, and fiber orientation were comparable, despite the differences in body size and, therefore, absolute muscle mass (Brown et al., 2010). Additionally, architectural studies of the female pelvic floor muscles showed that rats, compared to other commonly used laboratory animals, such as rabbit and mouse, were the closest to humans in terms of muscle design (Alperin et al., 2014). Moreover, the architectural difference index of rat pelvic floor muscles, which quantifies how closely rat muscle architecture resembles human muscle architecture, was comparable to the architectural difference index of the non-human primate (Brown et al., 2010; Lieber and Brown, 1992; Stewart et al., 2017). Furthermore, rat and human *in vivo* response to exercise shows similar qualitative and quantitative changes in plasma volume and in blood biochemical parameters (Goutianos et al., 2015). Additionally, the rat physiology is closer to human physiology, than mouse physiology is, making rats a widely employed preclinical model for toxicology and safety studies (Noto et al., 2018). Indeed, like in human, the rat genome contains genes involved in protein breakdown, and detection and detoxification of chemicals that have been lost in the mouse genome through natural selection (Gibbs et al., 2004). Finally, rats are 10-fold larger than mice, which impoves the ability to perform a wide variety of experimental procedures, collect larger samples, and study rare cell populations or low abundance molecules. The larger size of a rat also enables multiple concomitant measurements in a single animal, thus, reducing the number of research animals needed relative to a smaller mouse model.

Given that the rat model is widely used in multiple scientific fields and, in particular, in studies focused on skeletal muscles (Dwinell et al., 2011), we aimed to develop and validate an efficient and reliable protocol for MuSC isolation from rat skeletal muscles. The central role of MuSCs in the maintenance of muscle homeostasis and regeneration makes the ability to isolate and study MuSC autonomous properties of fundamental importance. Here, we describe for the first time an efficient and reliable method for isolation of rat MuSCs via fluorescent activated cell sorting (FACS) that relies on a single positive marker (VCAM-1) for identification of this cell population.

## RESULTS

### Determination of reliable positive markers for rat MuSC isolation

To isolate a pure MuSC population, we first identified commercially available antibodies that could be employed for the rat and tested them on cell preparations derived from the rat hind limb muscles to determine antibody binding specificity and optimal concentrations (Additional file 1: Figure S1). We selected the following positive markers: vascular cell adhesion molecule (VCAM-1, CD106); neural cell adhesion molecule (NCAM, CD56); and β1-integrin (CD29), based on the existing literature and compatibility with the rat. CD106, a transmembrane protein that belongs to the immunoglobulin superfamily, has been successfully used for isolation of mouse MuSCs (Liu et al., 2015). Importantly, expression of this protein in quiescent cells is required for maintenance of their basal function and prevention of premature lineage progression during cell activation (Choo et al., 2017). CD56 has been mainly described as a marker for identification of quiescent human MuSCs (Alexander et al., 2016; Castiglioni et al., 2014; Uezumi et al., 2016). CD29, has been previously used for the isolation of MuSCs from mouse, pig, and cow (Ding et al., 2018; Ding et al., 2017; Maesner et al., 2016). It is a member of the integrin family and interacts with both collagen and laminin depending on its heterodimer binding partner (Hynes, 2002). It is highly expressed in MuSCs and has been shown to be necessary for maintenance of quiescence in homeostatic conditions and cell proliferation after injury (Rozo et al., 2016).

As the first step, we titrated CD106, CD56, and CD29 antibodies through flow cytometry in order to confirm the expression of the related epitopes in the rat skeletal muscles and to determine optimal antibody concentrations. Hind limb muscles (tibialis anterior (TA), gastrocnemius (GAS), and quadriceps) were digested and pooled together. Analysis of the samples was performed on the LSR Fortessa where non-treated (NT) controls were compared to cells treated with one of five progressively increasing antibody concentrations (0.5, 1, 1.5, 2, 3 µg per 10^6^ cells) (Additional file 1: Figure S1a-S1c). Through calculation of the stain index (a measure of separation between the positive and negative populations) we determined the optimal concentration for all of the above antibodies to be 1.5 µg per 10^6^ cells. Similarly, we titrated antibodies for the negative selection markers: CD31, CD45, and CD11b. Each antibody was tested separately (Additional file 1: Figure S1d-S1f) to assess binding specificity and in combination to ensure that simultaneous use of multiple antibodies did not cause signal saturation (Additional file 1: Figure S1g). Optimal staining was achieved with 0.3 µg per 10^6^ cells for each antibody.

Guided by the previously established protocols in other animal models, we chose to test three isolation protocols using the TA muscle. Protocol 1 relied on CD106 as the only positive marker. Cells were concurrently stained for CD56 and CD29 or CD106 and CD29 in Protocols 2 and 3, respectively (Figure 1a). To set up the sorting protocol, we used single color and fluorescence minus one (FMO) controls to determine the gating system. Using forward and side scatter parameters independent of fluorescent signal, we first excluded cellular debris and cell clusters from the samples (Figure 1b-d, three top plots). For Protocol 1, we used DAPI negative staining to identify live cells. Within this live cell population, we then determined which lineage negative cells (CD31^-^/CD45^-^/CD11b^-^, Lin^-^) expressed CD106 (Figure 1b). Employing Protocol 1 we identified a single putative MuSC population (<u>P1</u>) that was further analyzed. For Protocols 2 and 3, we utilized the lack of negative selection markers and DAPI expression to identify live Lin^-^ cells, coupled with the expression of both positive markers to discern the presumed MuSC populations. Interestingly, expression of the CD56 marker employed in Protocol 2 was detectable in samples prepared from pooled hind limb muscle homogenate, but not in samples derived from TA alone, suggesting that the presence of CD56 protein might not be conserved among different muscles. We, therefore, excluded CD56 from further experiments. Despite the ellimination of CD56 antibody, we observed a clear separation of two cell populations based on CD29 expression in Protocol 2 (Figure 1c). We went on to further examine the following subpopulations, identified using Protocols 2 and 3: CD29^High^ (<u>P2</u>) and CD29^Low^ (<u>P2b</u>) for Protocol 2, and CD106^+^/CD29^+^ (<u>P3</u>) and CD106^-^/CD29^+^ (<u>P3b</u>) for Protocol 3 (Figure 1c-1d).

**Figure 1.**
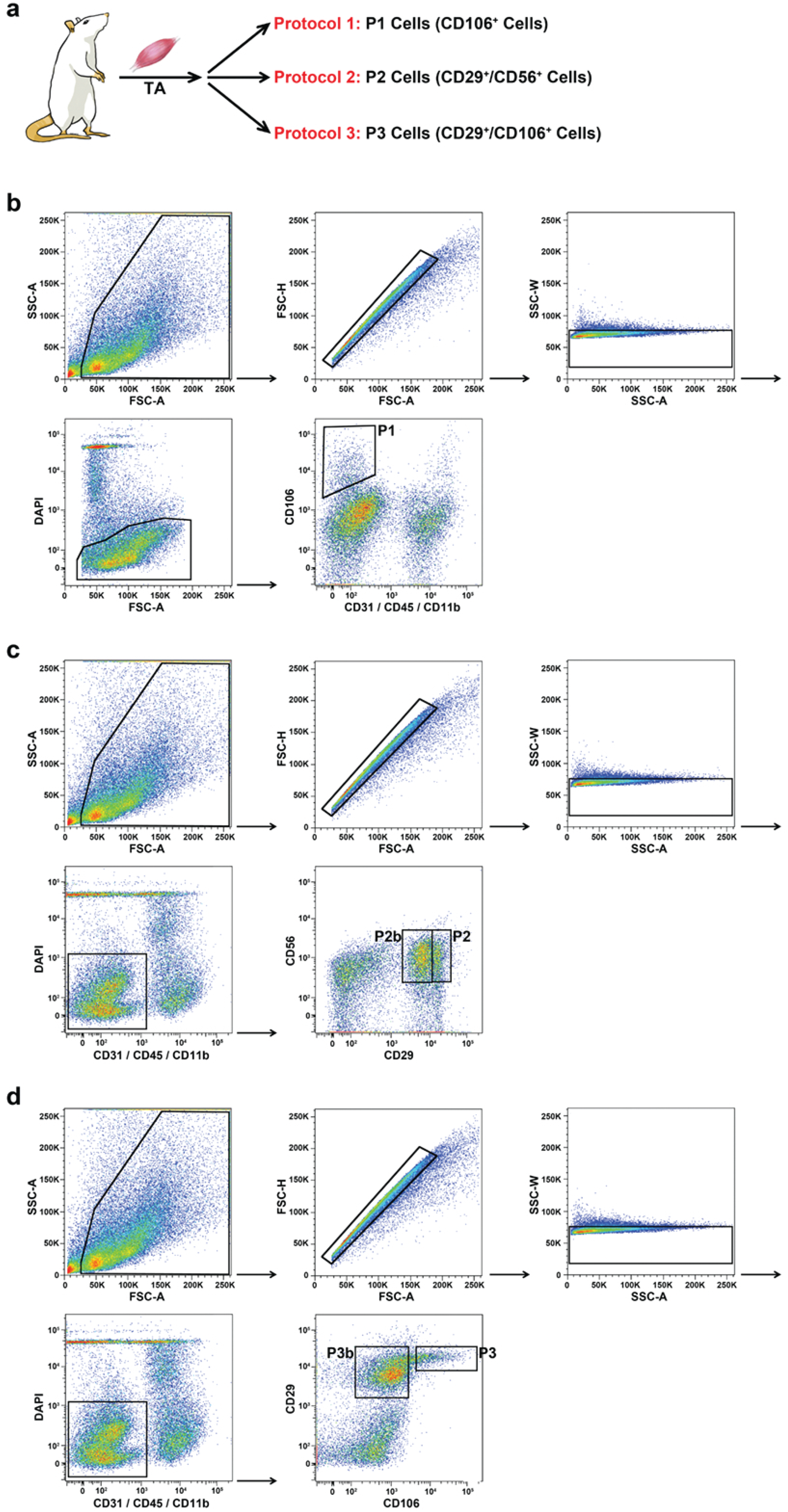
Gating approach for the three protocols employed for cell isolation. **(a)** Schematic of the experimental design for panels b-d. (TA: Tibialis Anterior). **(b)** FACS plots for the gating approach utilized in protocol 1. **(c)** FACS plots for the gating approach utilized in protocol 2. **(d)** FACS plots for the gating approach utilized in protocol 3.

### CD106 is a valid marker for isolation of rat MuSCs from tibialis anterior muscle

To test the identity and functionality of the five cell populations (P1, P2, P2b, P3, and P3b) described above, we isolated cells from TA muscle using BD Biosciences FACSAria II cell sorter. First, we evaluated the percentage of cells sorted using the three different protocols, with Protocol 2 limited to CD29 as the only positive marker (Additional file 2: Figure S2a). Protocols 1 and 3 yielded 1.6% of P1 (CD106^+^) and P3 (CD106^+^/CD29^+^) populations, which was significantly lower than 6.5% of putative MuSCs (P2 (CD29^High^) isolated employing modified Protocol 2 (Additional file 2: Figure S2a). P2b (CD29^Low^) constituted 9.3% and P3b (CD106^-^ /CD29^+^) 12.4% of the original sorted population (Additional file 2: Figure S2a). After isolation, cells were plated on laminin coated plates and either fixed either 2 or 12 hours later to assess cell identity or cultured in growth conditions for 3 and 5 days to determine their myogenic potential (Figure 2a). Expression of Pax7 in freshly isolated P1, P2, and P3 populations was around 90% and 80% at 2 and 12 hours after isolation, respectively (Figure 2b). In contrast, P2b (CD29^Low^) and P3b (CD106^-^/CD29^+^) populations expressed low levels of Pax7: 20% and 50% 2 hours after isolation, respectively (Additional file 2: Figure S2b). These results suggest that P1, P2, and P3 populations represent MuSCs, whereas, P2b and P3b likely do not.

**Figure 2.**
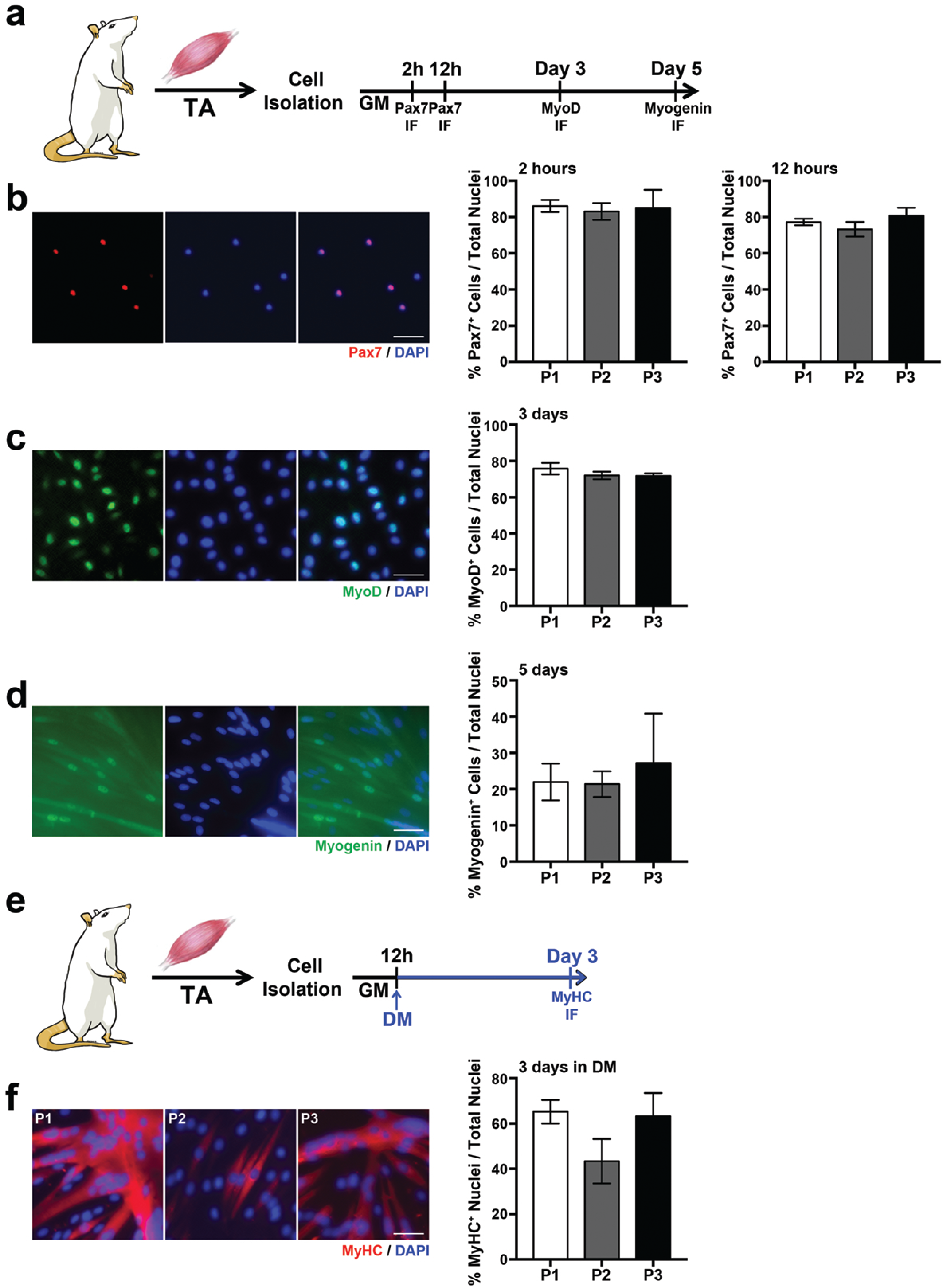
Phenotypic validation of cell populations isolated using three separate protocols. **(a)** Schematic of the experimental design for panels b-d (TA: Tibialis Anterior; IF: immunofluorescence; GM: Growth Media). **(b)** Representative immunofluorescent staining for Pax7 in freshly isolated cells on the left (scale bar: 50 µm); on the right, quantification of Pax7^+^ cells plated for 2 hours and 12 hours after isolation. **(c)** Representative immunofluorescent staining for MyoD in cultured cells on the left (scale bar: 50 µm); on the right, quantification of MyoD^+^ cells in culture for 72 hours. **(d)** Representative immunofluorescent staining for Myogenin in cultured cells on the left (scale bar: 50 µm); on the right, quantification of Myogenin^+^ cells in culture for 120 hours. **(e)** Schematic of the experimental design for panel f. (DM: Differentiation Media). **(f)** Representative immunofluorescent staining for myosin heavy chain (MyHC) in differentiated cells on the left (scale bar: 50 µm); on the right, quantification of MyHC ^+^ nuclei after 72 hours in differentiation media.

To assess the ability of the isolated cells to progress through myogenic lineage, we stained cells cultured in growth media for 3 and 5 days with antibodies against MyoD and myogenin, respectively. Around 80% of P1, P2, and P3 populations expressed MyoD after 3 days in culture, whereas, as expected, less than 40% of the P2b and P3b cells expressed MyoD (Additional file 2: Figure 2c and S2c). Moreover, 20% of P1, P2, and P3 cells expressed myogenin at 5 days, while myogenin expression was detected in less than 1% of P2b and P3b populations (Figure 2d and Additional file 2: Figure S2d). Taken together, these data show that only CD106^+^ (P1), CD29^High^ (P2), and CD106^+^/CD29^+^ (P3) cells express high levels of Pax7 and are capable of efficiently progressing through the myogenic lineage when grown *in vitro*.

To determine the ability of P1, P2, and P3 cell populations to complete myogenic differentiation, we placed the cells in serum-deprived media for 3 days, after which we assessed differentiation index by measuring the expression of myosin heavy chain (MyHC^+^), a terminal skeletal muscle differentiation marker, relative to the total number of nuclei (Figure 2e). P1 and P3 cells showed the highest differentiation index (over 60%) compared to 40% in the P2 population (Figure 2f). Moreover, P2 cells demonstrated a lower ability to fuse with each other, evidenced by sparse appearance of the myotubes relative to the P1 and P3 populations. (Additional file 2: Figure S2f). Consistent with their low ability to undergo myogenic commitment, the differentiation index of P2b and P3b cells was less than 2% (Additional file 2: Figure S2e). These results indicate that P1 and P3 populations have the highest differentiation and fusion potential. The above led us to conclude that both Protocols 1 and 3 accurately identify rat MuSCs capable of myogenic commitment and terminal differentiation. Despite high expression of Pax7, P2 cells (CD29^High^) isolated using Protocol 2, were not capable of efficient differentiation or fusion compared to P1 and P3 populations. We, therefore, excluded Protocol 2 from further analysis.

While Protocol 1 relies solely on CD106 as a positive isolation marker, Protocol 3 depends on two positive markers: CD106 and CD29. Importantly, cells obtained with either of these protocols did not differ phenotypically, indicating that both protocols yield comparable MuSC populations. Given fiscal and technical advantages of employing a single positive marker for the identification of MuSCs, we focused on validating Protocol 1 in five additional rat muscles.

### MuSCs can be efficiently isolated from a broad range of rat skeletal muscles employing CD106 as a positive marker

To validate our selected protocol, we tested its reliability and efficiency in isolating MuSCs from other rat skeletal muscles, specifically from GAS, a hind limb muscle commonly used in research, diaphragm (DIA), and the individual components of the pelvic floor muscle complex (coccygeus (C), iliocaudalis (ICa), and pubocaudalis (PCa)). To enhance population separation during the isolation process, and, therefore, increase MuSC yield, we initially focused on optimizing the gating system for Protocol 1 (Figure 3a). Based on MuSC size, determined during the initial gating system (Figure 1b), we first applied a gate to select for live small cells. Further gates were progressively designed to define single cell populations, Lin^-^ cells, and CD106^+^ cells. This new gating system was reproducible among all muscles evaluated, leading to a consistent isolation of 2 to 3% of cells from the total number of sorted cells and improving upon 1.6% yield of the previous gating system (Figure 3b, Additional file 2: Figure S2a and Additional file 3: Figure S3a).

**Figure3.**
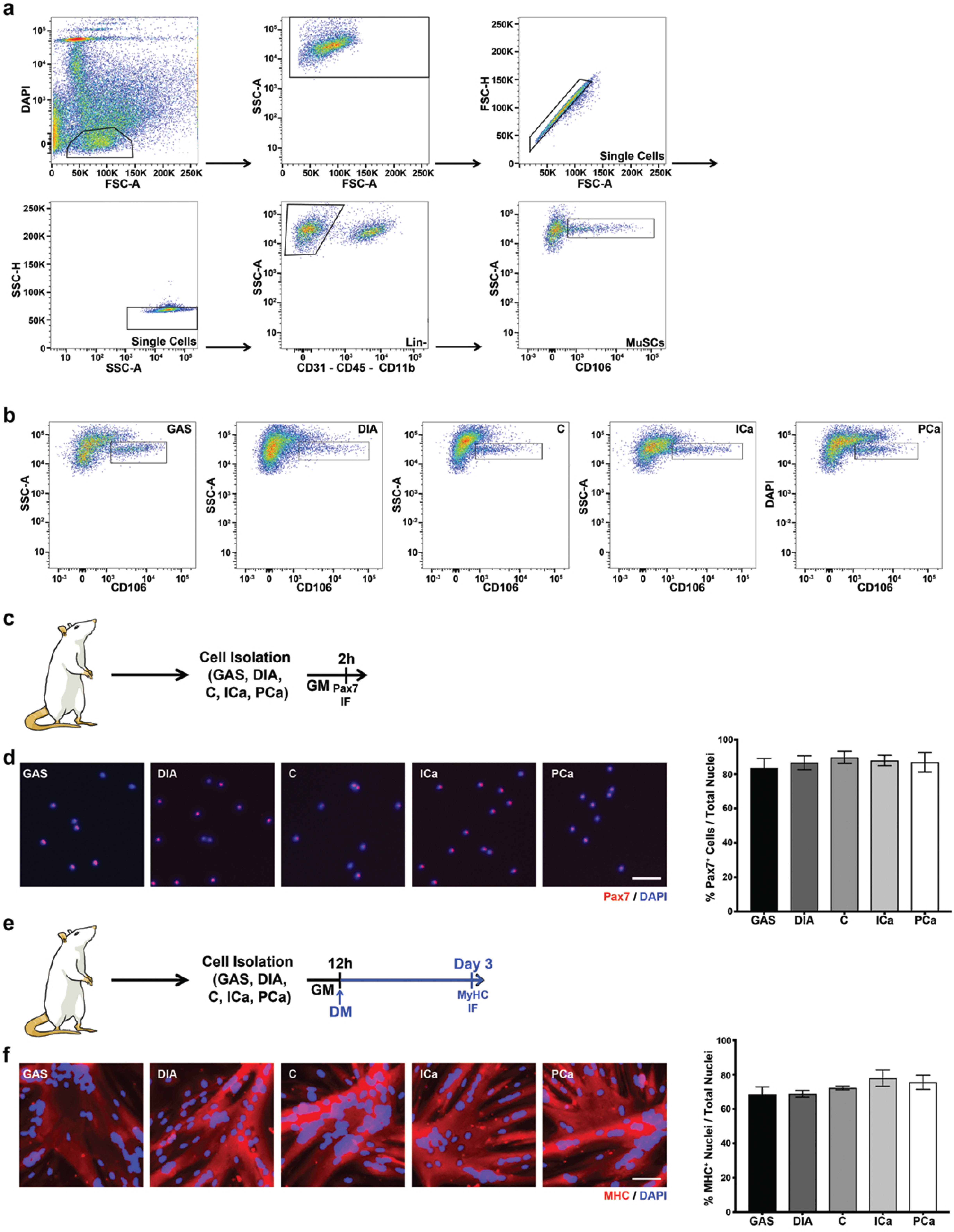
Phenotypic validation of cells isolated using Protocol 1 in gastrocnemius, diaphragm, and pelvic floor muscles. **(a)** FACS plots for the optimized gating system for protocol 1. (GAS: Gastrocnemius; DIA: Diaphragm; C: coccygeus; ICa: iliocaudalis; PCa: pubocaudalis; GM: Growth Media; IF: immunofluorescence). **(b)** FACS plots showing MuSCs (within the black gate) and in GAS, DIA, C, ICa, and PCa muscles. **(c)** Schematic of the experimental design for panel d. **(d)** Representative immunofluorescent staining for Pax7 in freshly isolated cells on the left (scale bar: 50 µm); on the right, quantification of Pax7^+^ cells plated for 2 hours after isolation. **(e)** Schematic of the experimental design for panel F. (DM: Differentiation Media). **(f)** Representative immunofluorescent staining for MyHC in differentiated cells on the left (scale bar: 50 µm); on the right, quantification of MyHC ^+^ nuclei after 72 hours in differentiation media.

Upon isolation, cells were plated in growth media for 2 hours, then fixed and stained with antibody against Pax7 to evaluate their identity (Figure 3c). Consistent with the results obtained for TA, around 90% of cells isolated from all other muscles using Protocol 1 expressed Pax7 (Figure 3d). Moreover, high expression of MyoD and myogenin was present when the cells were placed in culture for 3 and 5 days, respectively (Additional file 3: Figure S3b-S3e). When the cells were induced to differentiate in a serum-deprived media for 3 days, we observed 70% to 80% MHC^+^ nuclei (Figure 3e-3f). These results confirm that Protocol 1 can be reliably employed for isolation of a pure MuSC population, capable of myogenic commitment and terminal differentiation, across different muscle groups in the rat model.

## DISCUSSION

The current study demonstrates, for the first time, that CD106 (VCAM-1) is a reliable marker for the isolation of a pure MuSC population from various rat skeletal muscles. Indeed, isolated CD106^+^/CD45^-^/CD31^-^/CD11b^-^ cells express Pax7 at high levels (around 90%), and are capable of undergoing myogenic commitment, and full differentiation into myotubes *in vitro*.

Currently, the majority of MuSC studies are performed in a mouse, owning to the opportunities for genetic manipulation of this model (Huang et al., 2011). However, the small size of these animals significantly limits the amount of available muscle tissue, which in turn restricts the number of MuSCs that can be isolated. The above makes it hard to perform experiments demanding large cell numbers, such as RNA or ChIP sequencing. The insufficient yield of MuSCs from small mouse muscles necessitates pooling of different muscles from the same animal or pooling of the same muscle type from multiple animals to perform these molecular analyses (Sampath et al., 2018; van Velthoven et al., 2017). Pooling different specimens together masks the intrinsic variability of the different muscles or individual organisms potentially affecting results and their interpretation. Moreover, given recent evidence that MuSCs are highly heterogeneous, maintaining muscle and animal identity in future studies could enhance our understanding of diverse MuSC populations (Beauchamp et al., 2000; Cornelison and Wold, 1997; Kuang et al., 2007). Using the rat model, which is 10 times larger than a mouse (220g vs 25g for adult 8-week old animals), can help scientists circumvent these limitations and expand the existing studies to smaller muscles previously set aside due to technical constraints. For instance, the regenerative potential of MuSCs from extensor digitorum longus and soleus, small muscles that differ with respect to MuSC number and fiber phenotype, have never been directly compared (Soukup et al., 2002; Yin et al., 2013). The protocols employed for MuSC isolation from mouse muscles yield around 3% of the total number of sorted cells (Liu et al., 2015; Sacco et al., 2008). Using our optimized protocol, we obtain a similar proportion of the total sorted cells (Figure 3a and Additional file 3: Figure S3a). Given that the total number of MuSCs isolated from rat muscles is greater compared to mouse, utilizing the rat model precludes the need to pool samples from different muscles or multiple animals. This opens new avenues for investigations focused on the autonomous function of MuSCs derived from small-sized muscles, such as extensor digitorum longus and soleus. Larger animal models, such as non-human primates, pig, or cow, would, of course, allow isolation of an even greater number of MuSCs. However, these models are associated with significantly higher costs relative to the rat, and numerous constraints related to housing and handling of these species, as well as the need for specialized facilities.

In conclusion, a single positive selection marker can be used to reliably isolate MuSCs from a variety of rat skeletal muscles. The use of the rat model for the studies of MuSCs offers a major advantage to the skeletal muscle research field, enabling investigations of single muscles of different sizes and individual animals.

## MATERIALS AND METHODS

### Animals

Female 3 months old Sprague-Dawley rats were obtained from Envigo, Indianapolis, USA. Animals were euthanized using CO2 inhalation followed by bilateral thoracotomy, and hind limb muscles (tibialis anterior, gastrocnemius and quadriceps), diaphragm, and pelvic floor muscles (coccygeus, iliocaudalis, and pubocaudalis) were harvested. The University of California San Diego Institutional Animal Care and Use Committee approved all study procedures.

### Cell Isolation

Muscle stem cells (MuSCs) were isolated as described in Gromova et al., 2015 with minor revisions (Gromova et al., 2015). Tibialis anterior, gastrocnemius, diaphragm, iliocaudalis, pubocaudalis, and coccygeus were minced and subsequently incubated in 700 units/ml collagenase type II solution (catalog number: 17101-015, Life technologies, Gibco^®^) and collagenase and dispase II (catalog number: 04942078001, Roche^®^) solution (100 units/mL and 2 units/mL, respectively). Muscle tissue was then passed through a 10 ml syringe with 20 G needle and a 70 µm nylon filter. Antibody incubation was performed in 1 mL volume.

The positive markers used to identify the MuSC population were CD29 (catalog number: 102221, Biolegend), CD56 (catalog number: FAB7820P, R&D Systems), and CD106 (catalog number: 200403, Biolegend). The negative selection markers used to identify hematopoietic and endothelial cells were CD45 (catalog number: 565465, BD Biosciences), CD11b (catalog number: 562102, BD Biosciences), and CD31 (catalog number: NB100-64796AF647, Novus Antibodies titration was performed with LSR Fortessa. Determination of proper antibody concentration was achieved through calculation of the stain index (*FI*: fluorescent intensity; *SD*: standard deviation):

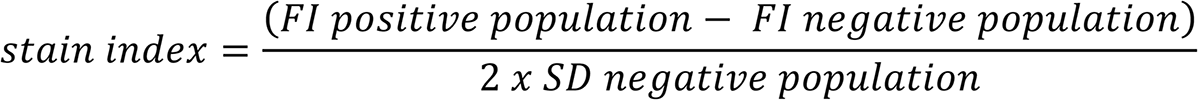

MuSCs were isolated with BD Biosciences FACSAria II cell sorter employing three different protocols: Protocol 1: CD45^-^ / CD11b^-^ / CD31^-^ / CD106^+^; Protocol 2: CD45^-^ / CD11b^-^ / CD31^-^ / CD56^+^ / CD29^+^; Protocol 3: CD45^-^ / CD11b^-^ / CD31^-^ / CD106^+^ / CD29^+^.

### Cell culture

After isolation, cells were plated (2500 cells per well) in laminin (catalog number: 11243217001, Roche^®^) coated plates in growth media (GM) (40% DMEM, 40% F10, 20% FBS, 1% Pen/Strep, 25 ng/mL βFGF). Cells were fixed at different time points after isolation (2 hours, 12 hours, 72 hours, 120 hours) to determine expression of myogenic markers. Myogenic differentiation was induced on 10,000 cells 12 hours after isolation, employing differentiation media (DM) (DMEM, 2% HS, 1% Pen/Strep). Differentiation was assessed 72 hours after media change.

### Immunostaining

Cultured cells were fixed with 4% paraformaldehyde, washed in PBS (phosphate-buffered saline), and incubated with blocking buffer before incubation with primary antibodies. Pax7 (1:100; catalog number: Pax7-c, DSHB), MyoD (1:100; catalog number: 554130, BD Biosciences), Myogenin (1:100; catalog number: 556358, BD Biosciences), and MyHC (1:100; catalog number: Mf20-c, DSHB) were incubated overnight in blocking buffer. Secondary antibodies (Alexa fluor 546 goat anti-mouse IgG and Alexa fluor 488 goat anti-mouse IgG, Thermo Fisher Scientific) were incubated at 1:250 dilution. Nuclei were identified with DAPI (1:1000; catalog number: 62248, Thermo Fisher Scientific).

### Imaging

Imaging was carried out using the Keyence BZX710 microscope. Image quantification was performed with Adobe Photoshop CS4 and ImageJ64. Images were not modified before performing cell counts.

## ACKNOWLEDGMENT

University of California San Diego Microscopy shared resources is supported by the NCI Cancer Center Support Grant P30 2P30CA023100-28 to UCSD Moores Cancer Center.

## AUTHORS’ CONTRIBUTIONS

FBS and MA contributed to study design, data analyses, and result interpretation. FBS, MW, and AC performed the experiments. FBS wrote the manuscript. MW, AC, and MA reviewed and edited the manuscript. All authors read and approved the final manuscript.

## FUNDING

The authors gratefully acknowledge funding by the National Institute of Health/ Eunice Kennedy Shriver National Institute of Child Health and Human Development, grants R01 HD092515 and R21HD094566, for the conduct of this research.

