## Supplementary Figures Legends for "Isolation of muscle stem cells from rat skeletal muscles"

**ADDITIONAL FILES LEGENDS**

**Additional file 1: Figure S1. Antibody titration for identification of optimal concentration for cells isolation. (a)** Schematic representation of the experimental design for panels B-H. **(b-h)** FACS plots for antibodies testing at five increasing concentrations. Results obtained for CD106 **(b)**, CD56 **(c)**, CD29 **(d)**, CD31 **(e)**, CD45 **(f)**, and CD11b **(g)** are represented. **(h)** FACS plots for combinatorial testing of all negative selection markers (CD31, CD45, and CD11b) at five increasing concentration. Each listed concentration refers to a single antibody.

**Additional file 2: Figure S2. Expression of myogenic markers in non-MuSCs isolated populations. (a)** Graphical representation of the % yield of isolated cells over the total number of sorted cells. Statistical test: one-way ANOVA. *: p-value<0.05; **: p-value<0.01; ***: p-value<0.001; ****: p-value<0.0001. **(b)** Quantification of Pax7^+^ cells plated for 2 hours and 12 hours after isolation. **(c)** Quantification of MyoD^+^ cells after 72 hours in culture. **(d)** Quantification of Myogenin^+^ cells after 120 hours in culture. **(e)** Quantification of MyHC^+^ nuclei after 3 days in differentiation media. **(f)** Representative low magnification immunofluorescent staining for MyHC in differentiated cells (scale bar: 100 μm).

**Additional file 3: Figure S3.** **Phenotypic validation of cells isolated using P1 in gastrocnemius, diaphragm, coccygeus, iliocaudalis, and pubocaudalis. (a)** Graphical representation of the percent yield of isolated cells over the total number of sorted cells. **(b)** Schematic representation of the experimental design for panels B and C. **(c)** Representative immunofluorescent staining for MyoD in cultured cells on the left (scale bar: 50 μm); on the right, quantification of MyoD^+^ cells in culture for 72 hours. **(d)** Representative immunofluorescent staining for Myogenin in cultured cells on the left (scale bar: 50 μm); on the right, quantification of Myogenin^+^ cells in culture for 120 hours.
