## Supplementary figures and images for "Isolation of muscle stem cells from rat skeletal muscles"

### Supplementary Figure 1

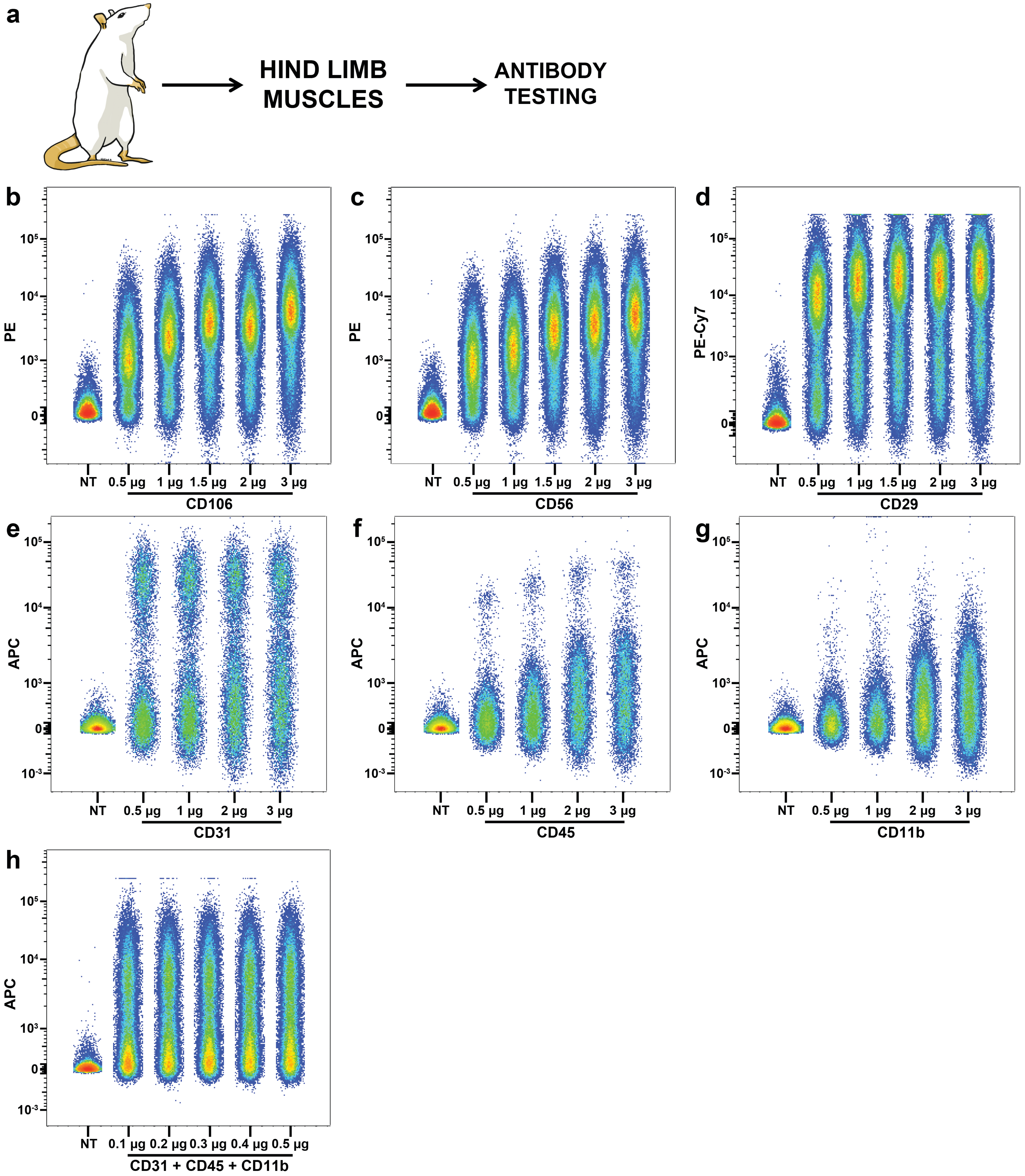

### Supplementary Figure 2

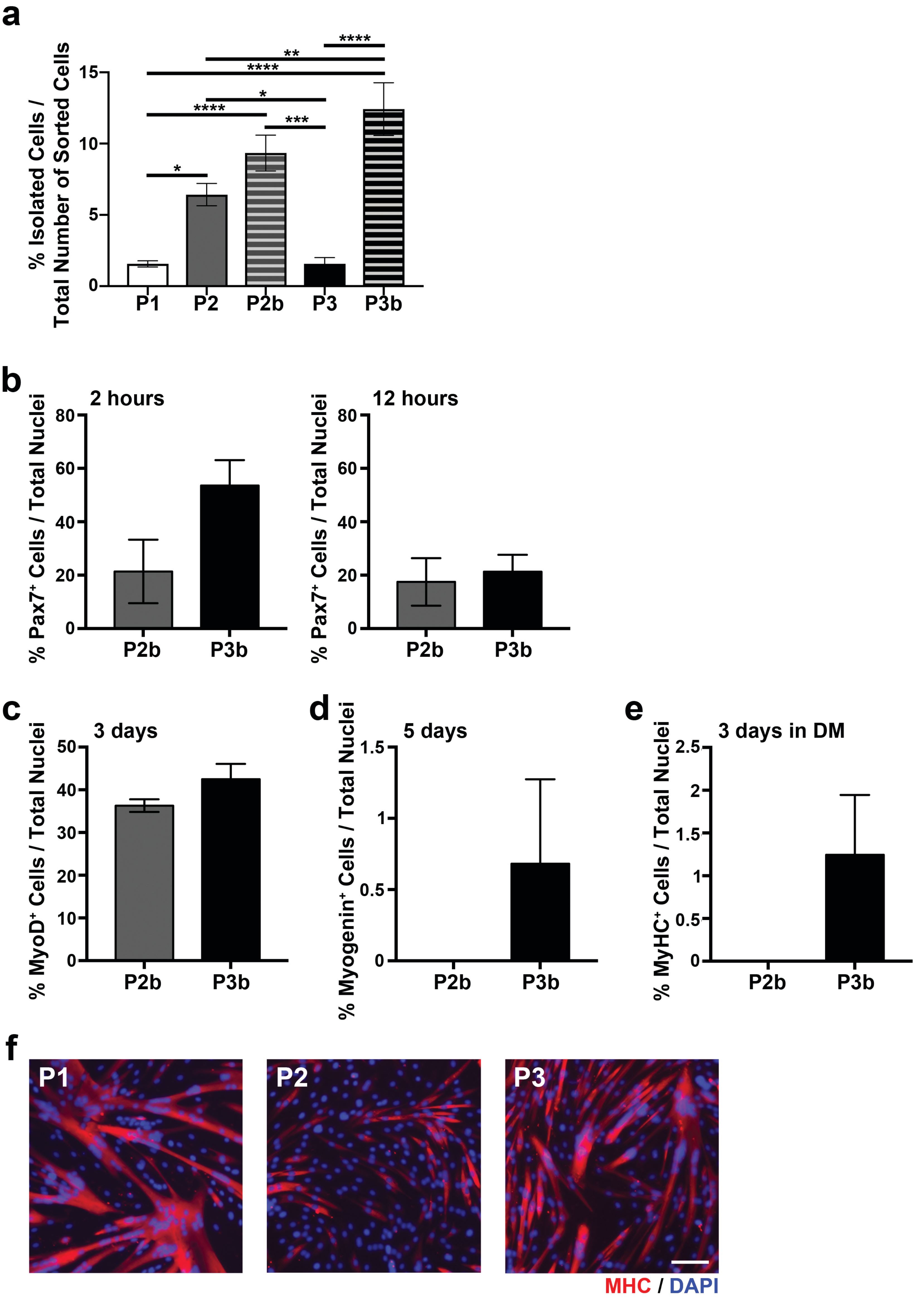

### Supplementary Figure 3

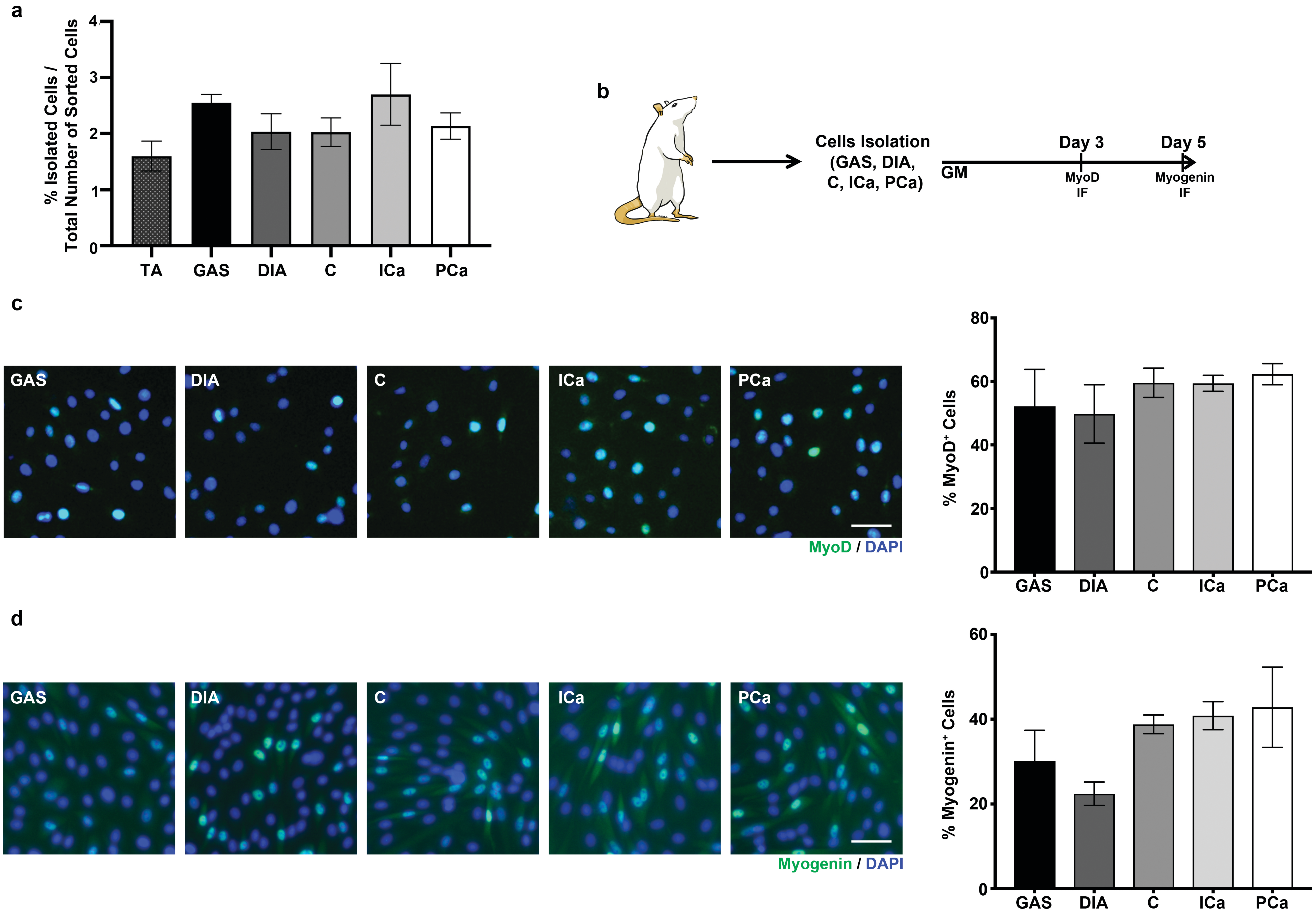
